# Membrane remodeling due to mixture of multiple types of curvature proteins

**DOI:** 10.1101/2021.10.02.462266

**Authors:** Gaurav Kumar, Anand Srivastava

## Abstract

We present an extension of the Monte Carlo based mesoscopic membrane model, where the membrane is represented as a dynamically triangulated surface and the proteins are modeled as anisotropic inclusions formulated as in-plane nematic field variables adhering to the deformable elastic sheet. In the extended model, we have augmented the Hamiltonian to study membrane deformation due to a mixture of multiple types of curvature generating proteins. This feature opens the door for understanding how multiple kinds of curvature-generating proteins may be working in a coordinated manner to induce desired membrane morphologies. For example, among other things, we study membrane deformations and tubulation due to a mixture of positive and negative curvature proteins as mimics of various proteins from BAR domain family. We also study the effect membrane anisotropy, which manifest as membrane localization and differential binding affinity of a given curvature protein, leading to insights into the tightly regulated cargo sorting and transport processes. Our simulation results show different morphology of deformed vesicles that depend on the curvatures and number of the participating proteins as well as on the protein-protein and membrane-proteins interactions.

**Significance:** Membrane remodeling requires highly orchestrated interactions between several types of lipids and curvature proteins. Experimentally probing the membrane deformation intermediates is non-trivial. For example, it is not known exactly how two or more different kinds of BAR domain proteins work in concert to induce and stabilize certain membrane curvatures. In this work, we use mesoscopic continuum modeling and explore the factors that induce and stabilize a range of membrane deformation due to a mixture of positive and negative curvature proteins.

## I. INTRODUCTION

Bio-molecular structures dynamically assemble, perform highly orchestrated biochemical functions, and disassemble seamlessly before reassembling to continue the cycle. Since the molecular structure and the biochemical pathways in biological processes are optimized over evolutionary timescales to perform elaborate functions, the “biological circuitry” is extremely sophisticated. Despite being under strong thermal noise and embedded in a crowded molecular milieu, the molecular machines and motors encrypt and process complex set of information in a precise manner. This information is embedded at multiple length and time scales that range from structures at atomic resolution to formation of molecular complexes and leading to subcellular and cellular processes such as ion transport across channels and pumps, fusion and fission in endocytic recycling, presynaptic neurotransmission, morphogenesis and metastasis to name a few.^1–3^

A paradigmatic example of this “biological circuitry” is the Clathrin-mediated endocytosis (CME) in vesicular trafficking. After five decades since its discovery in 1964 and countless seminal subsequent discoveries, it is now well established that more than 50 proteins make up this molecular machinery.^4–6^ Proteins, either individually or in complex are “called upon” from the cytosolic reservoir in a highly coordinated manner at different stages that include (i) initiation/identification of the endocytic site, (ii) recruitment of the cargo to the site, (iii) invagination/remodelling of the membrane, (iv) leakage-free scission (v) disassembly/uncoating of proteins and (vi) vesicle release for trafficking. These functions are carried out by disparate proteins that have been individually studied in great detail. However, we still lack a good understanding of how all these different components work together in a highly coordinated manner to drive vesicle formation. There are several examples like CME in the cell, which take place due to a large number of proteins and other biomolecules working as a ‘biological circuitry’.

BAR domain class of proteins are the quintessential curvature inducing scaffold proteins and are ubiquitously present in the endocytic recycling pathways and play a crucial role in vesicular transport processes in the cell.^5,7–12^ BAR family of proteins have a wide range of curvatures with proteins such as F-Bar and N-BAR exhibiting positive curvatures^5,13^ and proteins such as I-BAR exhibiting negative curvature.^14^ When these proteins bind to the membrane surface, they impose their local curvature on the membrane and also work in concert with each others at longer length scales to induce non-local curvatures on the membrane.^15–19^ Though the sub-cellular processes of vesicular transports is mediated by multiple curvature proteins interacting in concert with each other, this aspect has not been studied very much in reconstitution experiments.^20–24^ Also, lately role of flexible intrinsically disordered regions in scaffolding proteins has been shown to be important,^25–27^ which highlights the importance of considering the effective curvature of the scaffold protein as a distribution rather than having one fixed curvature value. Even the quintessential ‘‘scaffold” proteins such as BAR domain proteins can have extremely anisotropic shapes and a range of interaction strength with the membrane that significantly affects the sorting of proteins on the membrane surface.^28^ For example, BAR family protein Amphiphysin1 contains multiple disordered region^29^ and Epsin N-terminal homology domain^30^ as well as AP180 contains significant stretches of intrinsically disordered regions.^30^

The various experimental constraints as well as the resolution limits of imaging techniques makes it almost impossible to faithfully deconvolute the exact driving forces from the various components involved in membrane deformation. Molecular simulations can provide such insights but are formidable, even with low-resolution coarse-grained representation, due to the extremely large spatial and long temporal scales of the systems under considerations. Interestingly, mechanics-based mesoscopic models have provided important Physics-based insights into these emergent processes by capturing the membrane deformations observed in experiments. These mesoscopic methods are often developed using a multivariate optimisation framework where the final equilibrated membrane shape is derived by solving an energy-minimizing functional variational problem subject to problem-specific geometric constraints. Generally the membrane is presented as a discretized closed surface and proteins are implicit with the membrane or represented as inclusions.^31–34^ In particular, we point the readers to papers from John Ipsen and co-workers,^35–39^ Hiroshi Noguchi and co-workers,^40–42^ Thomas Weikl and co-workers,^43–46^ and the Mesoscopic Membrane with Proteins (MesM-P) model from Gregory Voth and co-workers.^47–49^ Due to its smooth particle applied mechanics (SPAM)^50,51^ based theoretically well-grounded framework, the MesM-P can provide phenomenological link between molecular interactions and macroscopic phenomena though the representation of particles is not in molecular detail and the membrane deformation studies are limited to one type of proteins.

Almost all of these work consider one protein type or a maximum of two proteins though these mesoscopic modeling work have provided important insights into the synergy between multiple scaffold proteins during membrane remodeling processes.^36,41,45,52,53^ For example, membrane remodeling due to two opposite curvatures of proteins was studied recently^54^ where the membrane is modelled as a triangulated surface and the proteins as arc-shaped particles. The model shows the membrane deformation due to two types of proteins with low or moderate scaffold curvatures. The method does not consider very high curvature proteins and also the protein-protein interaction is not considered in the work. Using a very different type of mesoscopic representation, Noguchi and co-workers have also recently explored the effects of two different types of proteins on membrane morphology. In their work, the proteins are modelled as “banana-shaped” chains of beads and the membrane is modelled as a two-dimensional sheet of beads. While studying the importance of membrane inclusions and scaffolding of proteins on the membrane tubulation,^40,53^ they show that if the curvature of inclusion and proteins both have the same sign, tubulation is promoted otherwise percolated-network is formed. When equal amounts of the two opposite inclusions are added, their effects cancel each other and in that case, tubulation is slowly accelerated. Also, in one of the first mesoscopic computational studies with more than one protein modulating membrane deformation, Ipsen and co-workers^36^ showed that membrane remodelling is driven by directional spontaneous curvatures of participating proteins. In their model, which we have borrowed and further extended in this work, the membrane is represented as a dynamically triangulated surface (DTS) and the proteins are modeled as a nematic field adhering to the deformable fluid membrane surface. In their exploration on multiple protein types, the study is limited to only two different types of proteins with no consideration protein-protein cross interactions for different type of proteins. Moreover, the model did not allow for variable surface coverage of proteins (all nodes on the triangulated surface were occupied by nematics/proteins). These studies are limited and quite few in numbers and a more extensive exploration in this area is needed.

Our nematic based modeling is the extension of the original method from Ipsen and co-workers,^35,37–39^ with an augmented Hamiltonian so that the method can be used to study membrane remodeling due to mixture of proteins with different curvatures working together with each other. In general, implicit field-based approach of most these mesoscopic modeling lends well to a conceptual understanding of physical factors leading to curvature formation but generally limits the scope of exploration in terms of heterogeneity in curvatures, anisotropy in membrane interactions and other ‘‘chemically specific” features including the variety in protein-protein interactions and surface localization of protein aggregates as well as in the study of assembly and scaffolding patterns of the participating proteins. We chose to extend the nematic-based model since the framework seamlessly allows us to incorporate the heterogeneity in the parameter space of the Hamiltonian in a mesoscopic setting. We can choose any curvature value and our results very clearly show that membrane deformation is driven not just by directional curvature of proteins but the interaction between the proteins and their surface coverage as well as the binding strength with the membrane play major roles in the remodeling process. We have also considered the interaction between same as well as different curvature proteins. A key aspect of the nematic membrane model is its ability to induce curvature even when the directional spontaneous curvatures are set to zero. We also show membrane remodelling due to a combination of straight and curved proteins as well as proteins that have different interactions with the membrane. To assure reliable interpretations of our results, we also rigorously test the results for convergence by starting from different initial membrane shape as well as randomized protein placements.

The rest of the paper is organized as following. We describe our model and algorithm in detail in the ‘Material and Method” section. In this section, we first describe the representation for membrane and protein and provide the formulations for the various interaction terms in the augmented Hamiltonian. We then discuss the dynamical triangulation Monte Carlo (MC) simulation algorithm that we implement for our model. Our work is rich in many use cases as we explore the full parametric space of effect of protein curvatures, protein-protein interactions, membrane-protein strength and surface coverage of the proteins on membrane deformation. We elucidate the full parameter space in the subsection titled “Parameters” in the ‘Material and Method” section. The “Results and Discussions” section follows next where we consider the many possibilities of the parameter space leading to formation of deformed membrane systems and we explore and report the salient design principles behind the different membrane shapes due to the various parameters. The paper ends with a short summarized conclusion.

## II. MATERIALS AND METHOD

Our model is an extension of the original model developed by John Ipsen and co-workers over the years,^35,37–39^ where the proteins are represented as nematics and fluid membrane is represented as a dynamically triangulated surface. The nematic lies on the local tangent plane of the vertex of the triangle. These nematics (proteins) are free to rotate in the local tangent plane and are denoted by 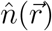. Here, the interactions between protein and membrane are modelled as anisotropic spontaneous curvatures of the membrane in the vicinity of the protein filament. Protein-protein interactions are modelled by the splay and bend terms of the Frank’s free energy for nematic liquid crystals. The total energy is the sum of membrane energy, energy due to interaction between protein and membrane and the protein-protein interaction energy. The model can also be seamlessly used to study the effects of pressure or surface tension on the membrane shape evolution by incorporating additional constraints in the Hamiltonian.

### A. Hamiltonian Model

#### 1. Membrane interactions

The energy of the bare membrane, which is the first of the total energy is written as:

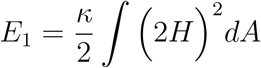

where *E_1_* is the Canham-Helfrich elastic energy for membranes,^55,56^ *κ* is bending rigidity and *H* is membrane mean curvature with *H* = (*c*_1_ + *c*_2_)/2. *c*_1_ and *c*_2_ are the local principal curvatures on the membrane surface along the orthogonal principal directions 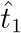 and 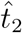.

#### 2. Protein – Membrane interactions

The coupling energy for the proteins and membrane interactions is written as *E*_2_ below.

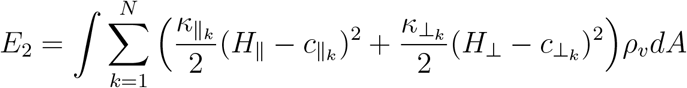

where *N* is the number of different curvature values of different types of proteins under consideration. *κ*_‖_ and *κ*_⊥_ are the induced membrane bending rigidities and *c*_‖_ and *c*_⊥_ are the intrinsic curvatures along 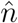 and 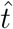, respectively. 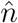 is the orientation of the protein and 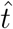 is its perpendicular direction in the local tangent plane. *H*_‖_ and *H*_⊥_ are the membrane curvatures in the direction of 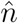 and 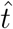, respectively and *H*_‖_ = *c*_1_ cos^2^*ϕ* + *c*_2_ sin^2^ *ϕ* while *H*_⊥_ = *c*_1_sin^2^ *ϕ* + *c*_2_ cos^2^ *ϕ*. *ϕ* is the angle between the direction of nematic orientation 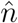 and the principal direction 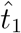. In the equation above, *k* in the summation indicates the number of different type of proteins, the *k* subscripts to *κ*_‖_ and *κ*_⊥_ indicate different different induced membrane bending rigidities and the *k* subscripts to *c*_‖_ and *c*_⊥_ are different principal curvatures of individual proteins. *ρ_v_* is the local coverage of nematic on the chosen vertex. If a nematic is present on a vertex, *ρ_v_* = 1, otherwise it takes a value of 0. This scheme allows us to have variable surface density of the proteins.

#### 3. Protein – Protein interactions

The protein-protein interactions is formulated as below:

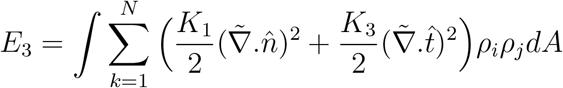

*E_3_* is the energy for the nematic-nematic interactions and is modeled from the Frank’s free energy for nematic liquid crystals. Here 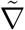 is the covariant derivative on the curved surface. *K*_1_ and *K*_3_ are the splay and bending elastic constants for the in plane nematic interactions. Here, a discrete form of this energy is used^35^ that makes the implementation amenable for the MC simulations and is expressed as:

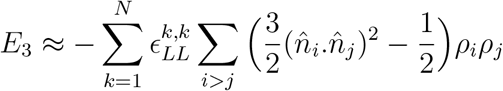

The above formulation is inspired from the Lebwohl-Lasher model.^57^ Here, *ε_LL_* is strength of the nematic interaction with a constant approximation (*K*_1_ = *K*_3_). The sum Σ_*i*>*j*_ is over all the nearest neighbour (*i*, *j*) vertices on the triangulated grid, promoting alignment among the neighbouring orientation vectors.

Besides the factors such as protein curvatures, protein-protein interactions and membrane protein interactions, the above Hamiltonian can also be augmented to study features such as conserved volume and surface tension effect. This is carried out by incorporating additional constraints in the Hamiltonian function. For example, effects of surface tension, which we report for one case later to highlight the scope of the model, can be studied by including an additional term *∫ σ.dA* in the Hamiltonian formulation.

### B. Dynamical triangulation Monte Carlo simulations

We use the MC optimization technique to solve the Hamiltonian and arrive at the equilibrium state of the deformed shape. The complete description of MC moves in the legacy model is detailed in the original work.^35,58^ We briefly describe it here for sake of completion and to avoid confusion that may ensue due to presence of multiple types of nematics. Four independent MC moves are required for the energy minimization to evolve the shape of dynamically triangulated surfaces with nematics and reach the minimum energy configuration. These MC moves are (i) vertex move, (ii) link flips, (iii) nematic shift, and (iv) nematic rotation. These MC moves provide the shape relaxation and fluidity in the system.

In the first MC move, a randomly chosen vertex *v* is displaced inside a cubical box, which is centered at the vertex and shown in Fig. 1(B). The move takes the membrane towards an equilibrium configuration. In the second MC move called the link flip, a set of two neighbor triangles is chosen and the common tether in between them is replaced by a new tether. This is created by a new connection in between two other non-connected vertices. Fig. 1(C) shows the link flip. In this case, the tether between *v* and *v*_2_ is replaced by creating new link between *v*_1_ and *v*_3_. This move guarantees that the vertex displacement is not controlled by the tether connection with its neighbors and thus ensures the fluidity in the system. The link flip move is known as the dynamically triangulated Monte Carlo (DTMC). The third MC move allows the nematic shift from one vertex to another. This move is shown in Fig. 1(D) where the nematic from vertex *v*_2_ is shifted to vertex *v*. Like the first move, this move too takes the membrane towards an equilibrium configuration. The fourth and final MC move allows the nematic rotation in the local tangent plane of the vertex. This move is shown in Fig. 1(E) where the nematic of vertex *v* is rotating in the local tangent plane of vertex *v*. This move is important for diffusion of proteins on the membrane surface as it affects protein assembly patterns and drives the membrane further towards the minimum energy shape.

**FIG. 1:**
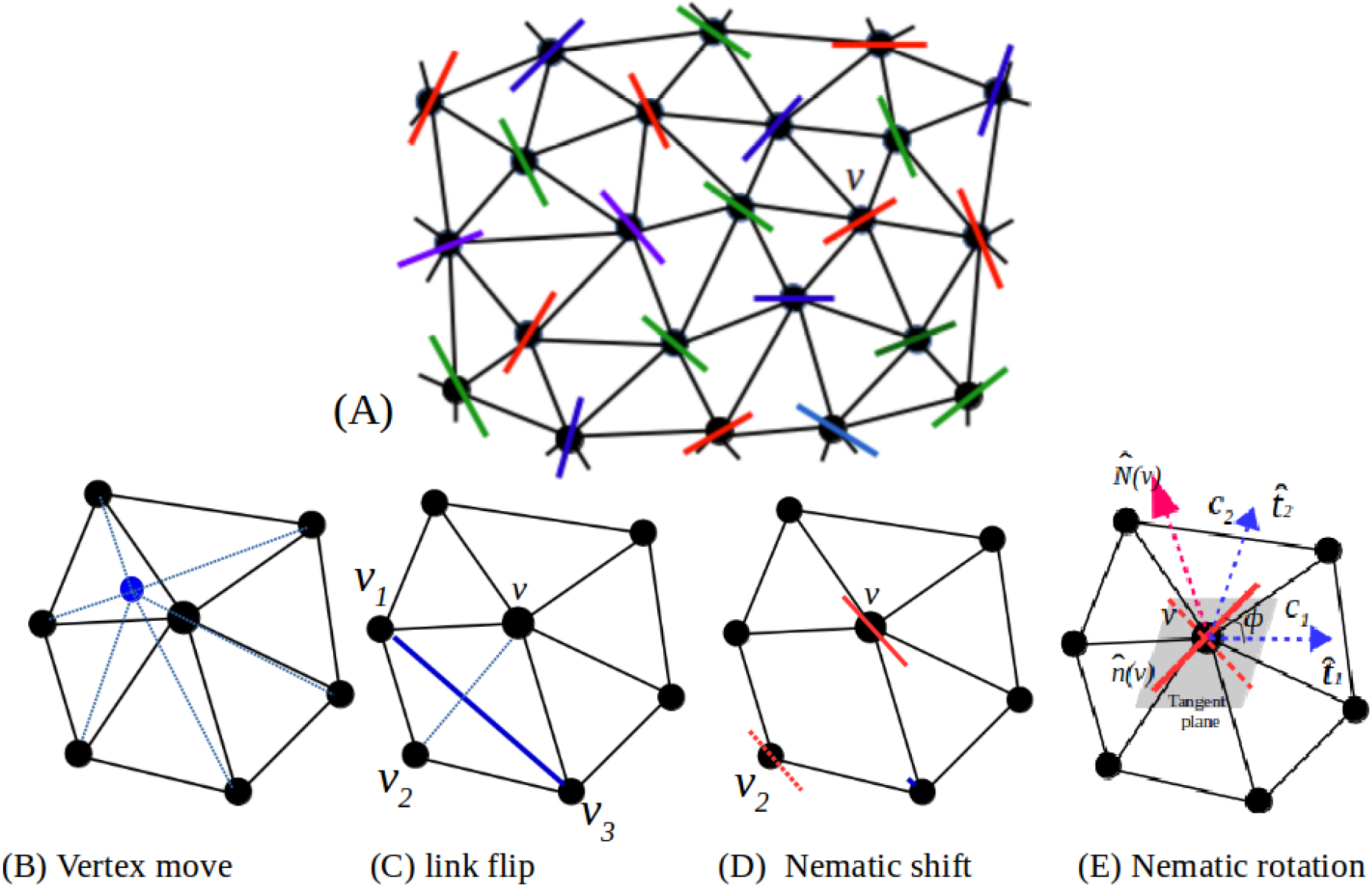
Schematics for the model. (A) The triangulated mesh represents the elastic membrane sheet and the vectors on the nodes represent the proteins as in-plane nematics. The model can account for multiple types of nematics presented as different colors in the schematics.(B-E) show the MC moves those are required for the energy minimization to evolve the shape of triangulated surfaces. (E) The various geometric parameters in the models, elaborated below, are represented here.

### C. Parameter space

Here, we elucidate the input parameters used in the simulation for multiple curvature proteins in form of matrices representation. If total number of different curvature proteins is *N*, then the nematic count is given as:

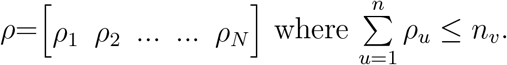

where *ρ*_1_, *ρ*_2_,…… *ρ_N_* are the number of different curvature proteins (nematics) and *n_v_* is the total numbers of the vertices on the triangulated surface. The total number of nematics cannot exceed the total number of vertices. Different proteins can have different curvatures, induced membrane bending rigidities and interactions.

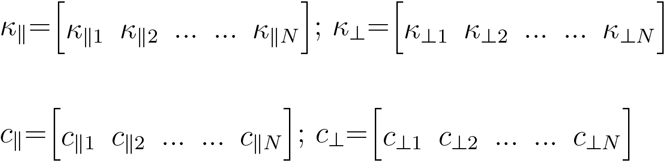

where *κ*_‖1_, *κ*_‖2_ and *c*_‖1_, *c*_‖2_……*c*_‖N_ are the induced membrane bending rigidities and curvatures of proteins along the direction of the nematics. *κ*_⊥1_, *κ*_⊥2_……*κ*_⊥N_ and *c*_⊥1_, *c*_⊥2_……*c*_⊥N_ are induced membrane bending rigidities and curvatures in the perpendicular direction of nematics.

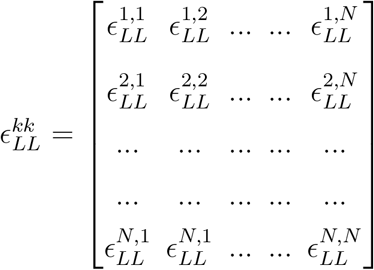

where 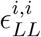 is the interaction between same curvature of nematics (same interaction) and 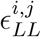 is the interaction between different curvature of nematics (cross interaction). We have considered both cases 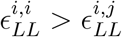 and 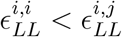 and show that how they affect the membrane deformation.

Our current implementation of the model is available on the Github repository https://github.com/codesrivastavalab/multiplescaffoldtypesDTMC/

## III. RESULT AND DISCUSSION

In this work, our primary focus is on the membrane remodeling due to mixture of different curvature proteins. Besides the curvature values, these proteins also have different membrane interactions. We also consider cases where a single protein can exist with variable curvatures due to fluctuations in the flexible regions of the proteins and we present such protein systems as a distribution of different curvatures with same membrane interaction strength. In all our simulations, proteins are randomly distributed on the spherical vesicle in the initial setup and their organization and membrane deformation evolve and converge to the equilibrium solution. But when needed, to test for convergence and to avoid possibilities of kinetically-trapped metastable states, we also carry out additional simulations with different initial shapes of the membrane. In the results below, we look into the protein organization and final vesicle shape due to interplay between proteins and membrane and we explore the factors leading to tubulation and membrane deformation that depend of non-local emergent factors such as bundling tendencies in proteins as well as curvature mediated protein organization. Before we move to salient findings of our studies, we present some simple cases that highlights the power of the method in studying curvature-proteins mediated membrane deformation in a high throughput manner.

Fig. 2 shows the vesicle deformation due to two different proteins with equal but opposite curvatures ±0.5 and with all vertices on the triangulated sheets occupied. In the Fig. 2, the x-axis shows that the number of positively curved proteins decreases from 90% to 10% while the number of negatively curved proteins increases from 10% to 90%. Protein-protein interaction is shown on the y-axis. We first discuss the two extreme examples on the left bottom and top right of Fig. 2. In the bottom left, the shape evolves to a discocyte to accommodate the curvature propensities due to 90% positively curved proteins and only 10% negatively curved proteins. The initial shape of the vesicle does seem to put a constraint on the final evolved shape, which is clearly evident by the oblate shape of the right top example in Fig. 2. In the oblate shape, the high density negative curvature nematics are accommodated by local dimples and flatter surfaces. Since the protein-membrane interaction (imposed membrane rigidity *κ*_‖_) is also low in this example, so curvatures are not imposed strongly. Another instructive use case is the middle example in Fig. 2 (±50% with *ε*_‖_ = 3.5). The positive nematics (Blue) localize the top and bottom ends where the membrane curvature is outwards and the negative nematics (Red) populate the regions where the surface has negative curvature. This propensity is also visible in other examples in Fig. 2 and the final deformed shape is optimized based on the composition and interaction strengths.

**FIG. 2:**
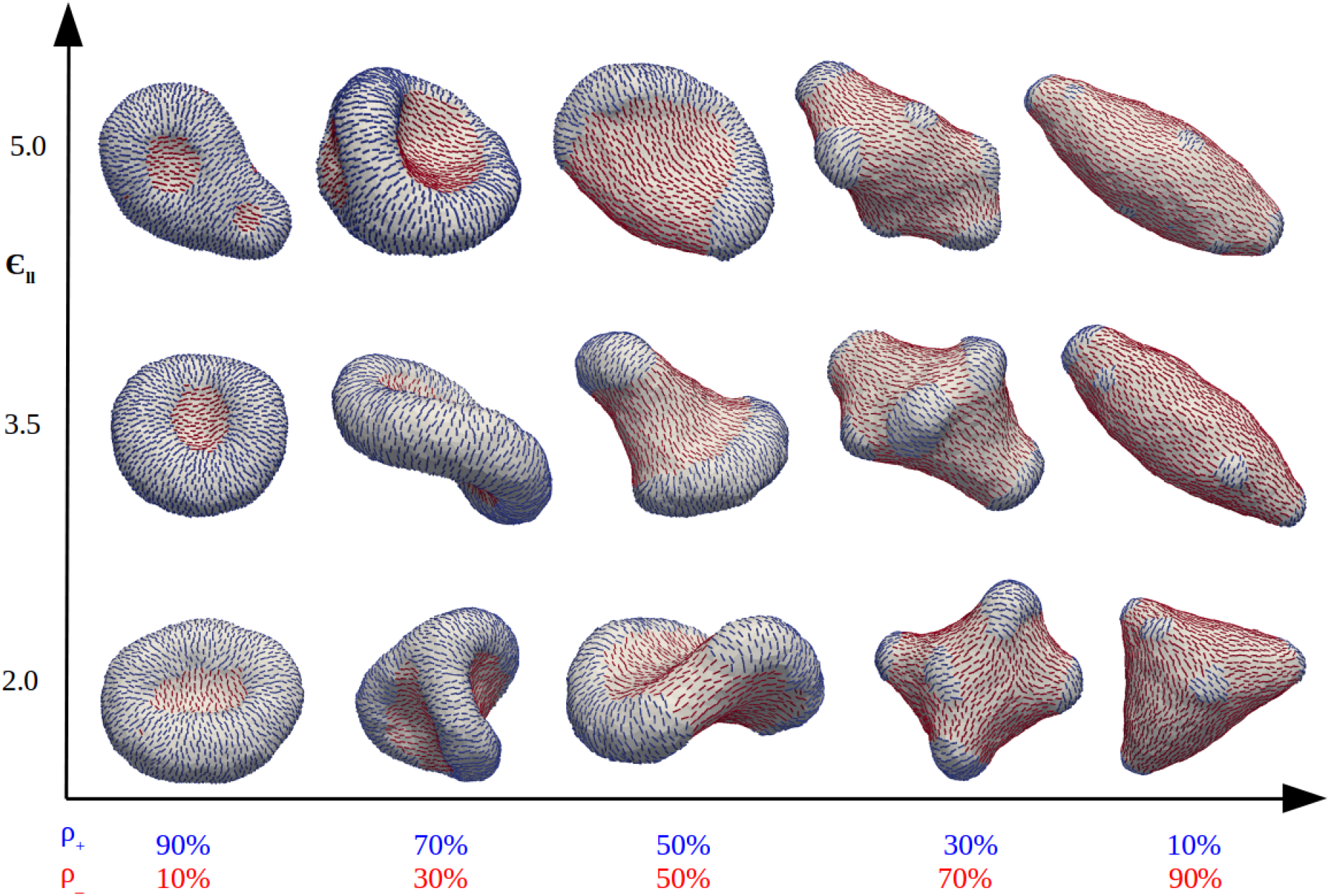
Panel shows the vesicle deformation due to two different proteins with curvature (±0.5). Here, different shapes are obtained with different protein numbers and different protein-protein interactions. On the X axis, number of positively curved protein is decreasing while negatively curved proteins in increasing. On the Y axis, nematic-nematic interaction 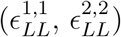 is increasing. Other parameters are *κ* = 20, *κ*_‖_ = 5 in *κ_B_T* unit.

In an another example highlighting the scope of this model, we show simulation results for the membrane remodeling due to two, three, four and five different curvature proteins. Left panel of Fig. 3 shows membrane deformation due to (a) two, (b) three, (c) four and (d) five different proteins with curvature values of (±0.5), (0, ±0.5), (±0.5, ±0.8) and (0, ±0.5, ±0.8), respectively. Here, the total protein coverage is 100% and populations are equally divided into different curvature proteins. In these examples, we have taken bending rigidity of the membrane as *κ* = 20*k_B_T*, induced membrane bending rigidity as *κ*_‖_ = 5*k_B_T* and nematicnematic interaction as 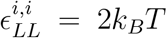. We discuss the details of Fig. 3(a-d) in the first subsection below and focus on how interactions between protein types drive their demixing and induce non-local large membrane deformation. In the right panel of Fig. 3, we highlight how differential expression of proteins of different curvature affects membrane deformation. This figure shows the vesicle deformation results due to three different curvature proteins where x-axis represents the different curvature value of proteins under different compositions while y-axis represents protein-protein interaction. In these results, the curvature of the three participating proteins are 0.3, 0.4 and 0.5 and the membrane-protein interaction *κ*_‖_ is 10 *k_B_T*. This example highlights the effect of up regulation or down regulation of certain protein on the overall membrane deformation. For example, all other parameters remaining same, different final shapes are formed when the relative composition of the proteins are changes (horizontal rows). Also, for the same composition, the deformation profiles are different when protein-protein cross interactions are different. Together, these examples highlight the capability of our model towards studying membrane remodeling due to mixture of multiple types of curvature proteins.

**FIG. 3:**
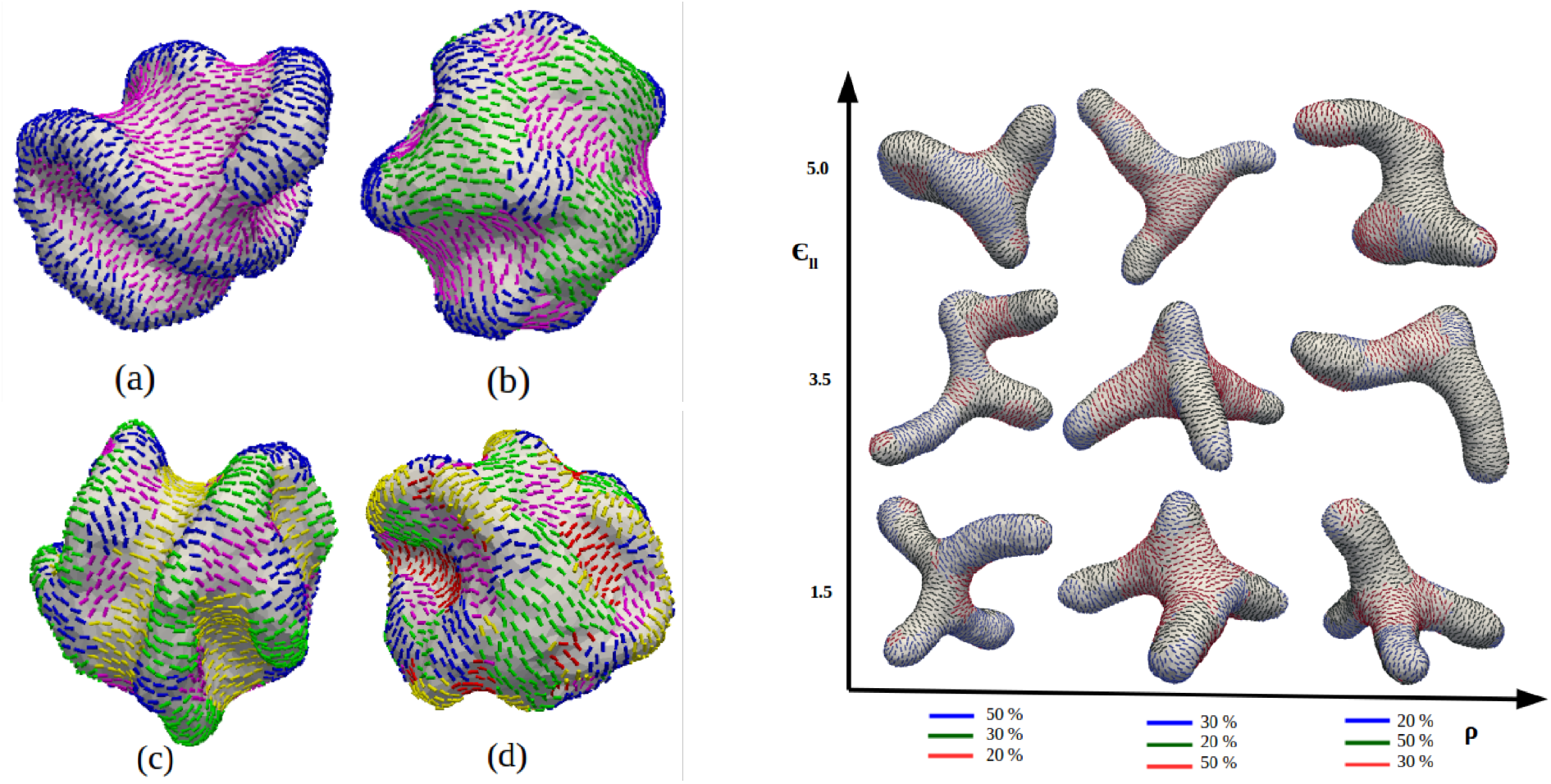
Deformed vesicle morphologies obtained by Monte Carlo simulation. (Left panel) Results for two, three, four and five different curvatures of proteins with curvature values (±0.5), (0, ±0.5), (±0.5, ±0.8) and (0, ±0.5, ±0.8). Different colors show the different values of protein’s curvature. Other parameters are *κ* = 20, *κ*_‖_ = 5 and 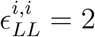 in *k_B_T* unit. (Right panel) Vesicle deformation due to three different curvature value of proteins under different compositions and protein-protein interactions 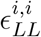. Other parameters are *κ* = 20, *κ*_‖_ = 10 in *k_B_T* unit.

In general, we start all our systems with a spherical vesicle and with random protein distribution for all use cases. However, we also carry out multiple simulations with different initial shape to ensure convergence. We find that if the energy and other features such as volume and area show convergent behaviour consistently for extended steps then we obtain converged solution of shapes. For example, in the Fig. 4, we simulate a system with two types of nematics having equal curvatures (0.6) and existing in ratio of 3:2. Only half the vertices are populated with nematics, which is equivalent to 50% membrane surface coverage. In this example where we highlight our convergence prescription, we have taken bending rigidity of the membrane as *κ* = 20*k_B_T*, induced membrane bending rigidity as *κ*_‖_ =10*k_B_T* and nematic-nematic interactions as 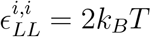 and 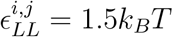.

**FIG. 4:**
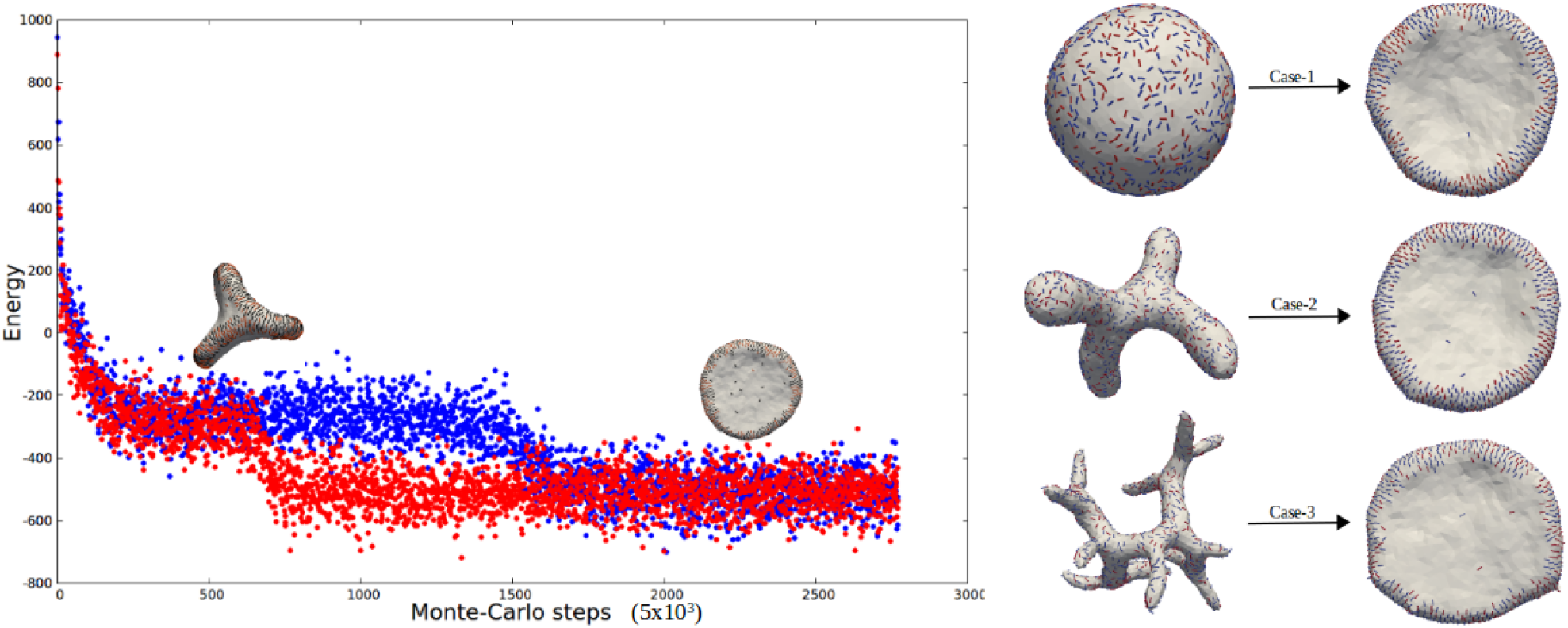
(Left panel) The minimized energy convergence is a necessary criterion to show that the deformed structure in not a metastable structure. (Right panel) Disc formation with three different starting shapes leading to identical final shapes

We do find that sometimes metastable states (such as the branched shapes) were formed instead of the final disc shape as seen in the left panel of Fig. 4. If the run was not extended, the branched shape could be misinterpreted as the final converged shape. However, on energy analysis, we noticed that the branched shape has higher energy as compared to the disc shape. We show the energy comparison in the left panel of attached Fig. 4 for the branched shape (in Blue) and disc shape (in Red). It is important to run the MC simulations for an extended period of time to facilitate better convergence. In most cases, issues related to non-transient “locked” intermediate shapes are addressed by running longer simulations. However, we are aware and do acknowledge that there are still possibilities of insurmountable kinetic traps occurring in the evolution of some of the use cases. And simply running the simulations for longer may not always lead to a final converged shape. For a more involved convergence analysis, we also test for volume and surface area convergence and we often rerun the simulations with different initial shape and protein distributions. For example, in the example discussed here, we ran the simulations with three distinct initial membrane shape as shown in the right panel of Fig. 4. To truly randomize the initial shape, we make sure that these non-spherical initial shapes do not occur in the pathway of the simulation where sphere was the initial shape. Ideally, a full “free-energy” convergence analysis should be done with advanced sampling schemes to guarantee true convergence. We are fully aware of the limitations of testing convergence with our current prescriptions and are working on an advanced sampling based approach to guarantee the convergence in situations where a metastable state may exist in a deep free-energy basin. We intend to address this aspect appropriately soon. In the following, we put forward some interesting design principles that emerge from our parametric studies using the developed model.

### A. Weak interactions between protein types drive their demixing and induce non-local large membrane deformation

In cellular context, two or more different proteins frequently interact with each other during membrane remodeling processes. ^20–23^ However, due to the spatial and temporal resolution of the imaging techniques in membrane remodelling experimental studies, it is non-trivial to clearly observe the interplay between the proteins leading to membrane deformation. In this section, we show some simulation results where two different proteins having opposite curvatures come together for membrane deformation. We find that this does not always lead to membrane deformation. In particular, we focus on the factors that determine the localization and assembly patterns of the participating proteins leading to deformation. Fig. 5 shows the simulation results for two different curvature of proteins with different numbers.

**FIG. 5:**
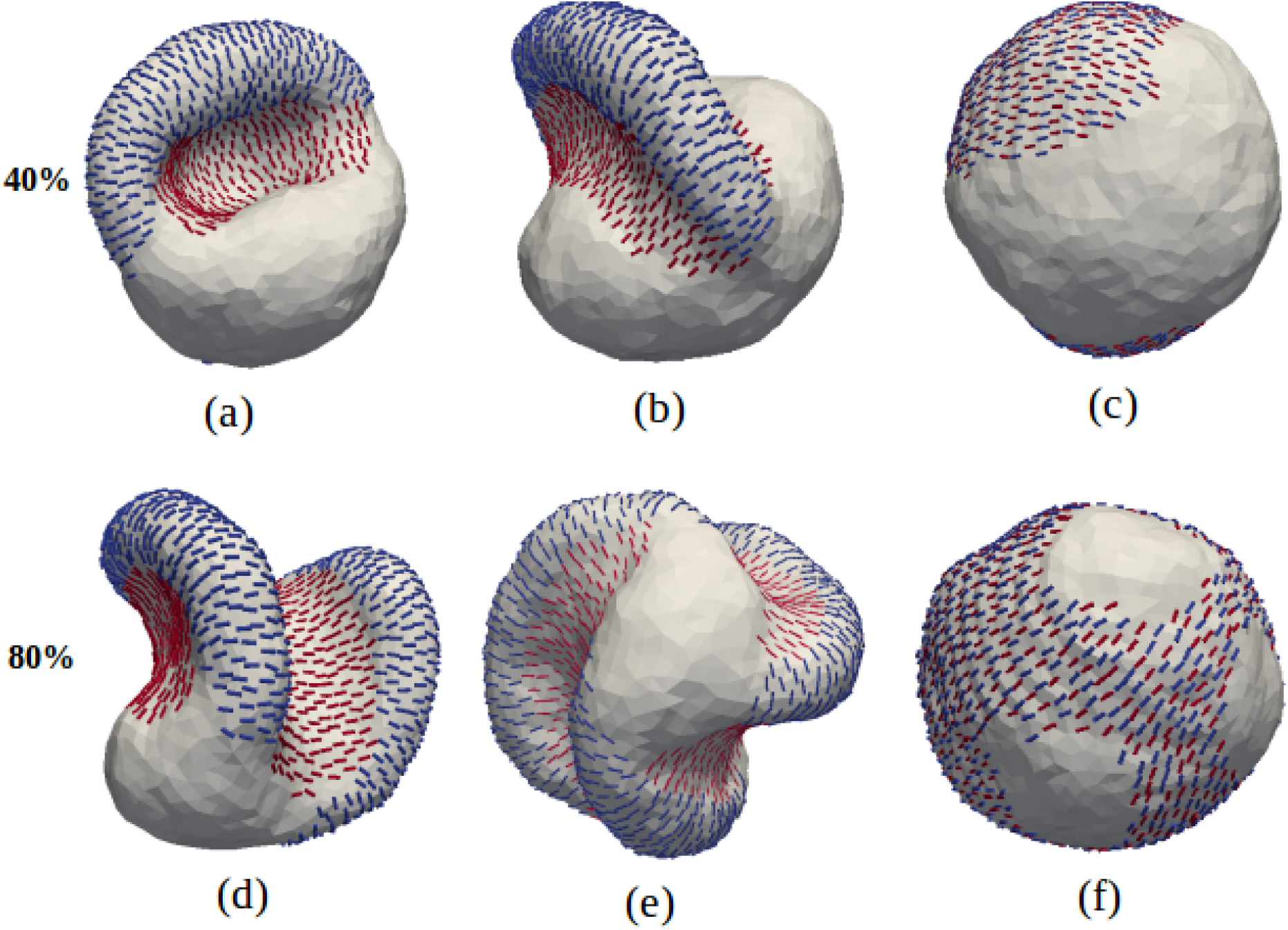
Vesicle deformation results with non zero interactions between the two different curvature of proteins. Here protein’s number is fixed 40% for upper panel and 80% for lower panel. For (a,d) 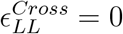 (b,e) 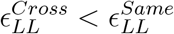 (c,f) 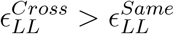. Other parameters are *κ* = 20, *κ*_‖_ = 10 and 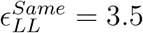 in *k_B_T* unit.

Here the interaction between the different curvature proteins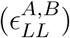 is also included and denoted by 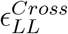. Protein coverage in upper and lower panels are 40% and 80%, respectively. Here the curvature of the two different proteins are ±0.5. Proteins with positive curvature are shown by Blue color while the negatively curved proteins are shown by Red color. Fig. 5(a,d) shows the simulation result with 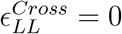, which indicate that there are no interactions between the Red proteins and the Blue proteins. It is worth noting that the two types of proteins assemble as separate demixed groups in close proximity to each other. This is because positively curved proteins generate a negatively curved regions in the areas adjacent to their local assembly and it becomes energetically most favorable for the negatively curved proteins to migrate in that region and stabilize the overall deformation profile. This is a clear example of curvature induced localization and assembly of second protein type. We also observe similar behavior in the left panel of Fig. 3(a-d) where the proteins stabilize the final deformation by localizing in preferred curvature regions. We use the results in Fig. 3b to accentuate this point again. Three different proteins with different curvatures are used in Fig. 3b. Blue and Pink nematics represent +0.5 and −0.5 curvatures, respectively while the Green represents 0.0 curvature. As is clear from the figure, the final deformed membrane has Blue nematics localizing on the positive outwards curvatures while the Pink nematics populate the negative inwards curved part of the deformed vesicle. The Green nemantics, with zero curvature, are localized in the curvature regions between the other two.

In Fig. 5(b,e), we keep the cross-interaction term 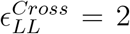 such that 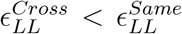. As a result, the two protein are allowed to interact with each other but due to higher self-interaction, they tend to be demixed and localize in preferred curvature regions as in the first case. We also see in Fig.2 that different final shapes are obtained due to different value of nematic-nematic interaction and different proteins number. It is worth noting that with increasing interactions, proteins with higher number tend to assemble in one place rather than be localized in different pockets. This also leads to larger scale deformation rather than punctuated localized deformation. In systems with two proteins having equal and opposite curvatures, we find that a strong protein-protein interactions lead to complete mixing of the two kinds of proteins. In Fig. 5(c,f), we modeled this situation where 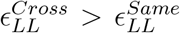. Due to the higher cross interactions, we observe that the two oppositely curvature proteins completely mix with each other. The fully randomly mixed arrangement of oppositely curved proteins in close proximity to each other has a compensatory effect on the curvature formation and no shape deformation is visible.

Our simulations also provide some very interesting insights into cases with two different proteins having same curvature but different induced membrane bending rigidity. This is important for many classes of curvature proteins including the various nexin complexes (SNXs) that have similar scaffold curvature but with specialized membrane adaptor domains such as PH domain and PX domain with preference for certain organelle membranes^5,59–61^. These proteins bind with different strength to a given membrane despite the same scaffold curvatures. Fig. 6 shows the membrane remodeling with two different proteins shown by Blue (A) and Red (B). We fixed the membrane-protein interaction of protein A to 10 *k_B_T* and varied the induced membrane bending rigidity of protein B, which is shown on the y-axis. The numbers of both proteins are changing on the x-axis but the total count is fixed and equal to 50% of the vertices. Here, the protein-protein interaction between same types of proteins is 2 *k_B_T* and between different types of proteins is 1.5 *k_B_T*. We observe that as the induced membrane rigidity (or membrane interaction strength) of second type of protein with same curvature increases, there is a higher propensity to form tubules. In the top row, tubes are formed with as less as 30% of protein B while that is not the case with weaker membrane interactions. What we also found very interesting in that there is complete mixing between the proteins as they prefer the same curvature, which suggest that assembly and localization of curvature proteins in not just affected directly by how strongly they bind to the fluid membrane but also by the strength of the protein-protein interactions. The bottom row of Fig. 6 have same results across the composition, which is reassuring to see since the differentiating parameter (membrane interaction strength) is same for both types of proteins.

**FIG. 6:**
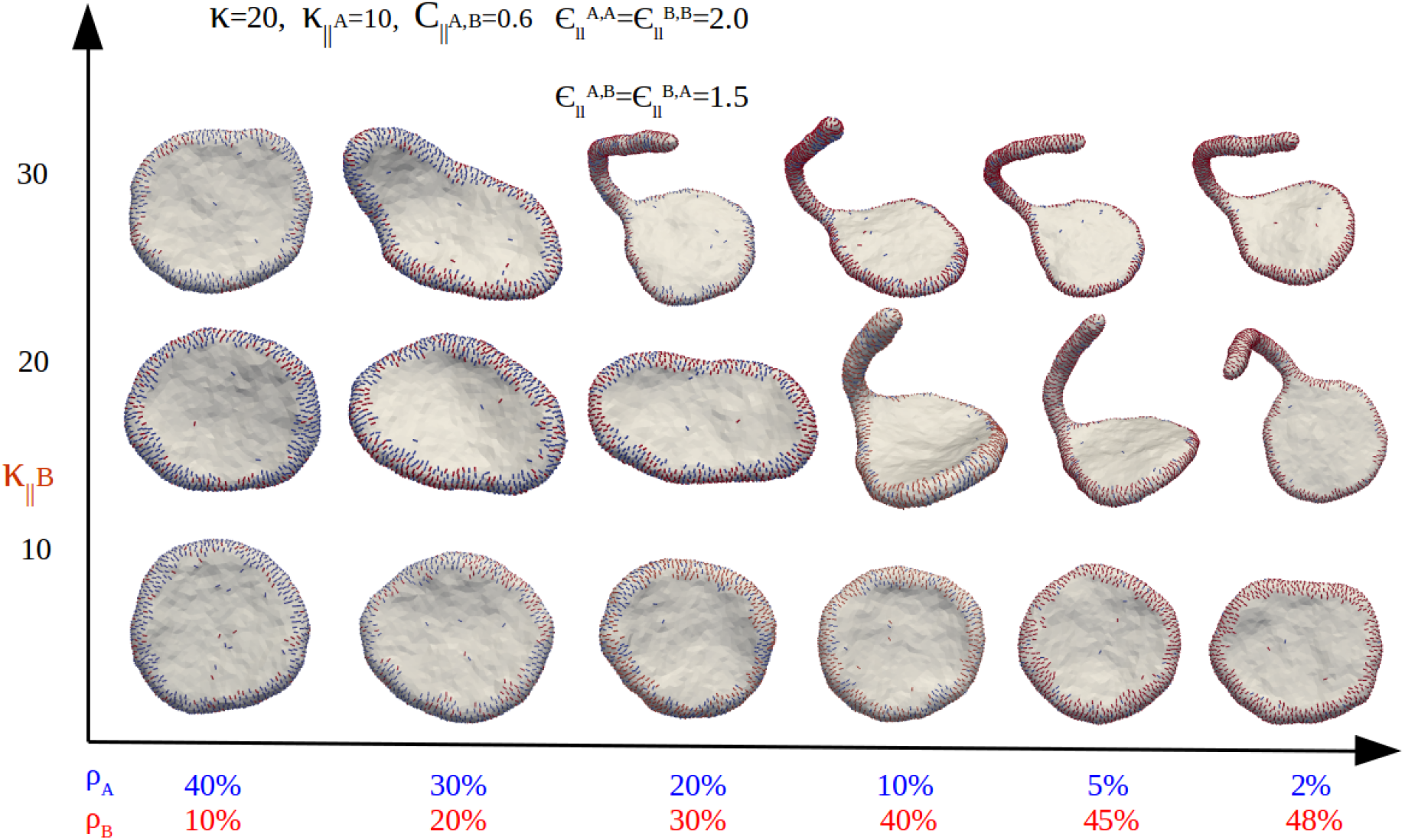
Vesicle deformation due to two different proteins with equal curvature *c*_‖_ = 0.6 and different induced membrane bending rigidity. Other parameters are *κ* = 20, 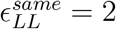 and 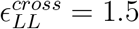 in *K_b_T* unit.

In the next section, we show the importance of protein-membrane interaction towards curvature formation with special focus towards tubulation propensities.

### B. Reticulation is facilitated by high membrane-protein interactions

Our analyses show that the strength of induced membrane bending rigidity is the primary driving force for tubulation to occur. For example, in Fig.7, we present results for the vesicle deformation due to proteins with three different curvatures. Here protein curvatures are 0.5, 0.75 and 1.0 with a count of 30%, 40% and 30%, respectively. X axis shows increasing membrane protein binding interactions while the Y axis show an increasing protein protein interaction. As can be seen from Fig.7, different shapes are obtained due to different values of protein membrane interaction and protein protein interaction. We observe that the reticulaion propensity increases with the induced membrane bending rigidity (membrane interaction strength). For example, we see highly reticulated structures with thinner tubes for higher induced membrane bending rigidity. This behavior is consistent across all protein-protein interactions, which suggests that though protein-protein interactions drive localization and certain degree of deformation, a high induced membrane bending rigidity is required to form highly tubulated profiles.

**FIG. 7:**
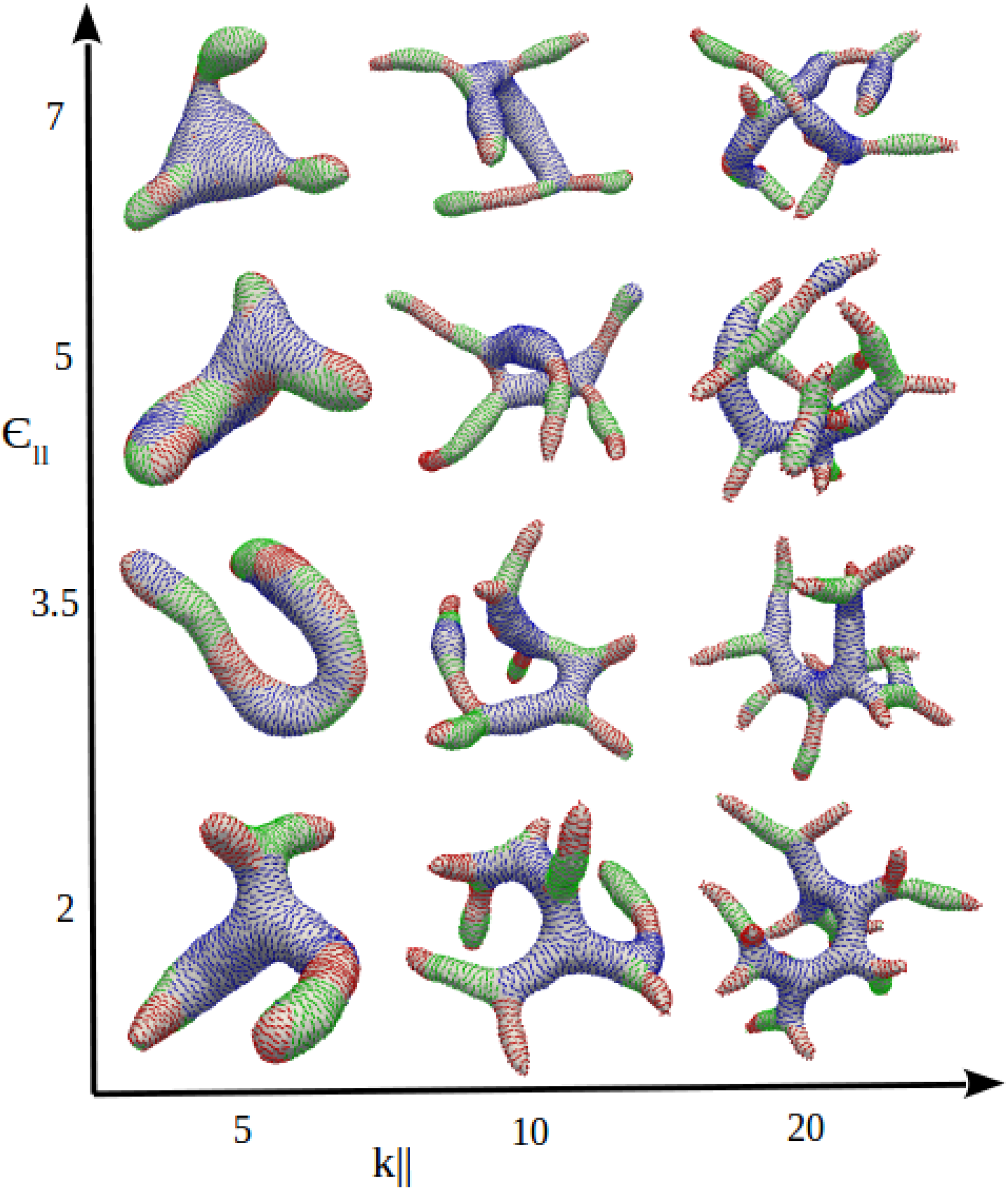
Vesicle deformation due to proteins with three different curvature values 0.5, 0.75 and 1.0 with protein coverage 30%, 40% and 30%. Different shapes are obtained due to different values of membrane nematic interaction and nematic-nematic interaction 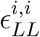.

To highlight the dominant role of membrane-protein interaction over protein-protein interactions during tube generation process, we were able to create a case where tubes are formed from vesicle with two different proteins that are not interacting with each other at all. These results are presented in Fig. 8, where the total protein count is equal to 50% of the surface coverage but the ratio of protein count is different in the four cases. The induced bending rigidity for tightly bound protein (shown in Red) and loosely bound proteins (shown in Blue) are 40 *k_B_T* and 20 *k_B_T*, respectively. All other parameters are same as for the system shown in Fig. 6.

**FIG. 8:**
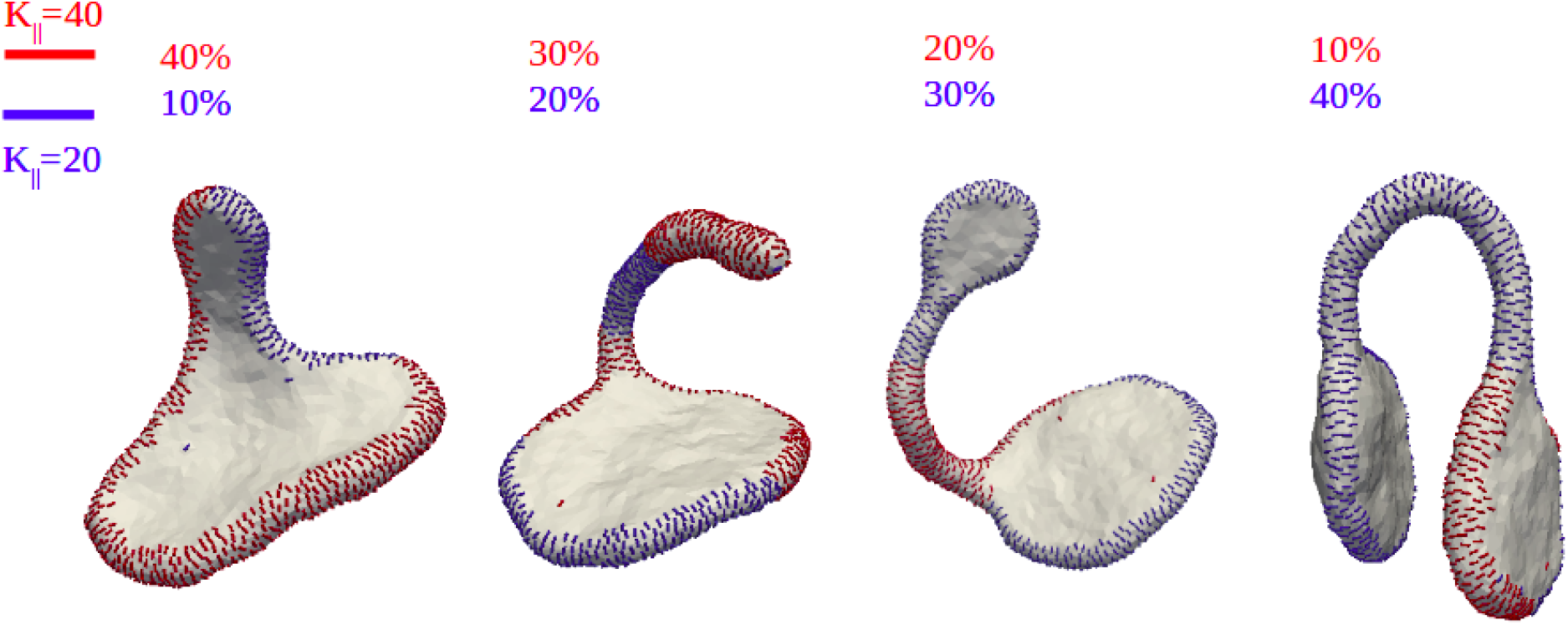
Vesicle deformation due to two different proteins with equal curvature *c*_‖_ = 0.6 and different induced membrane bending rigidity. Other parameters are *κ* = 20, 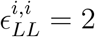 in *k_B_T* unit.

Interestingly, we also find that it takes lesser number of proteins to form tubes or large deformation when multiple types of scaffold protein as opposed to a single type of scaffold proteins. In Fig. 9, we show membrane remodeling due to one type of scaffold proteins. Here all parameters are same as Fig. 6. The protein number is varying on the x-axis and the induced membrane bending rigidity is varying on the y-axis. Around 70% proteins coverage is required for the tubulation but we find in Fig. 6 that when two different proteins interact with the membrane, lesser overall protein count can cause tubulation. The finding that less proteins are required for membrane remodeling when two different proteins interact the membrane with different induced membrane bending rigidity is quite interesting. We believe that presence of variety of curvature proteins allows the system to stabilize the evolving heterogeneous curvatures that occurs during the membrane remodeling process.

**FIG. 9:**
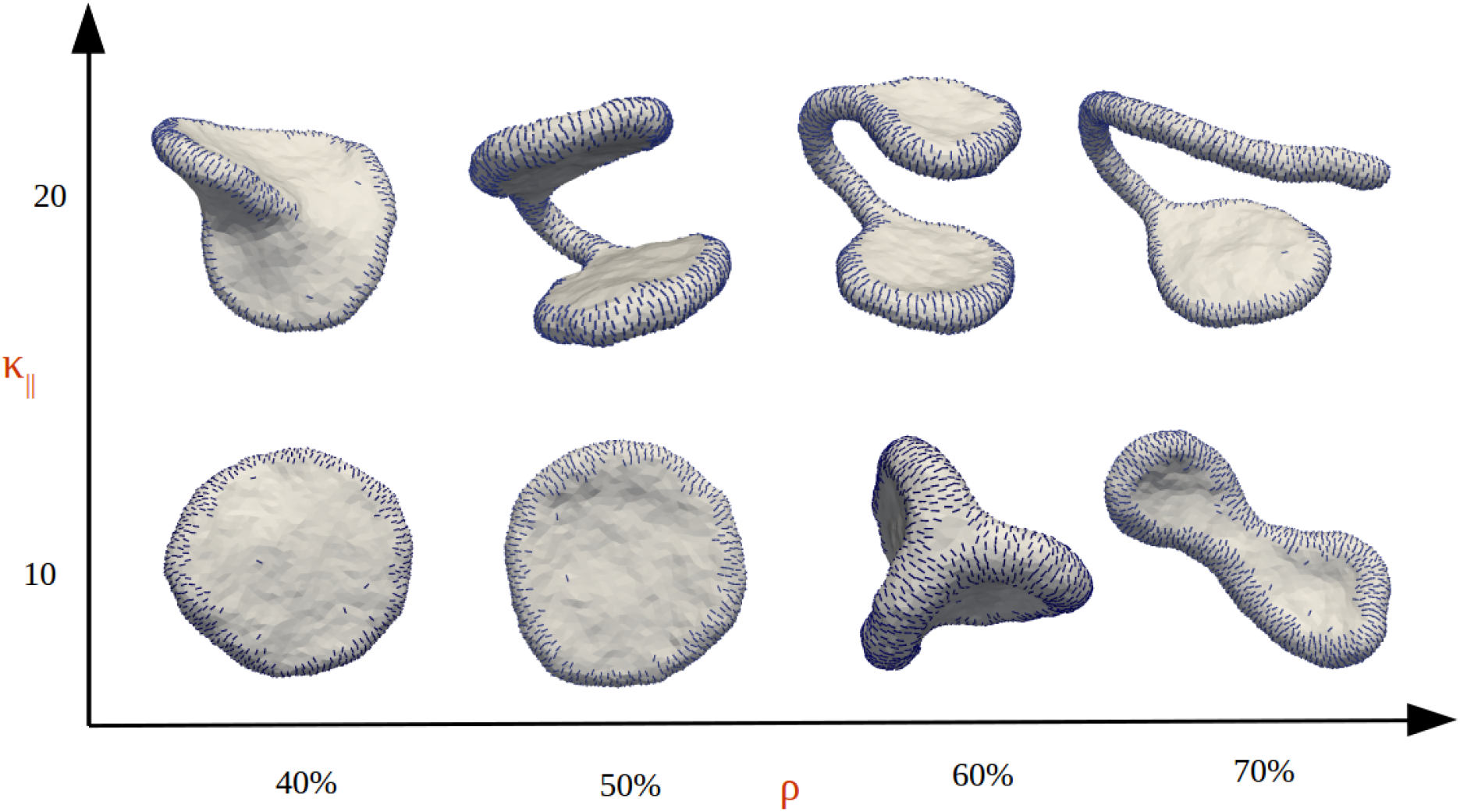
Vesicle deformation due to one type of proteins with curvature *c*_‖_ = 0.6 and Other parameters are *κ* = 20, *κ*_‖_ = 10 and *ε_LL_* = 2 in *k_B_T* unit.

Endoplasmic reticulum (ER) like morphology is another example where it is known that mixed curvature protein systems is thermodynamically favorable to induce and maintain very thin reticulated structures with multiple complex junctions between tubules. In ER, tubes are stabilized by Reticulon and Reep proteins while the junction between them is stabilized by protein Lunapark. Fig. 10a shows the membrane tubulation with two different proteins with curvature 0.5 and −0.3. Positive curvature proteins form tube while the negative curvature proteins form the junction between the tubes. Here the protein number for positively and negatively curved proteins are 90% and 10% respectively. Other parameters are *κ* = 20, *κ*_‖_ = 15and 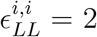 in *k_B_T* unit. In Fig. 10b, we show the membrane tubulation for three different curvature proteins with a distinct triple junction, a topology feature commonly observed in ER tubes. In heavily reticulated systems such as those belonging to ER and Golgi Apparatus, it is energetically more favorable to have a combination of positively and negatively curved scaffold proteins to stabilize a highly heterogeneous topology. Presence of one kind of scaffold protein leads to certain constraints that disallows morphologies like triple junction and neck regions where both positive and negative curvatures coexist and requires proteins with opposite curvature to stabilize such essential mixed curvature zones.

**FIG. 10:**
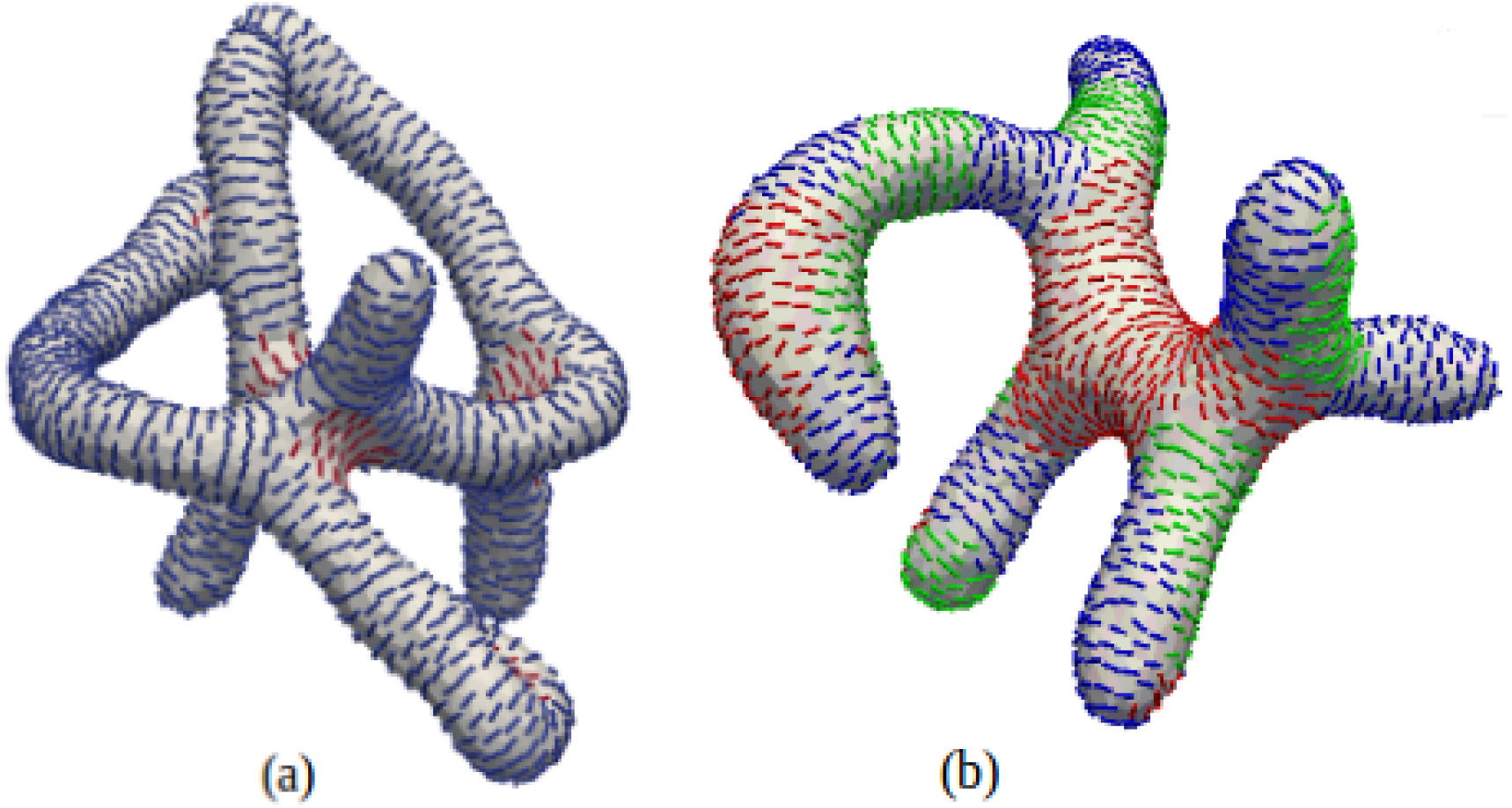
(a) Membrane tubulation due to two different proteins with curvature 0.5 and −0.3. Positively curved proteins are stabilized on tube while the negatively curved proteins are stabilized at the junction between tubes. (b) Membrane tubulation due to proteins with variable curvature. Here the curvatures are 0.3, 0.4 and 0.5. Other parameters are *κ* = 20, *κ*_‖_ = 15and 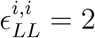 in *k_B_T* unit.

### C. Flat proteins with zero curvature can cause tubulations due to bundling tendencies

Membrane tubes are quite common in cell functions and are formed by the curved proteins and bundling tendency of the proteins. Tube formation due to bundling tendency of protein is briefly discussed in ref^39^. In this work, we show membrane tubulation due to combination of two different curvature proteins. Fig.11a shows the membrane tubulation due to two curved proteins with curvature *c*_‖_ = 1.0 and −0.05. The tubes localize the positively curved proteins (Blue) while the negatively curved proteins localize in the valley region (Red). Proteins that don’t have curvature but can form bundles like microtubule are also capable of creating membrane tubes. Fig.11b shows the membrane tubulation due to bundling of the proteins. Here one protein is has zero curvature (Red) *c*_⊥_ = 1.0, *c*_‖_ = 0 and another one have a positive curvature (Blue) *c*_‖_ = −0.05. Arrangement of the protein on the tube is different in Fig.11a and b. In Fig.11a, proteins arrange themselves in the transverse direction to the length of the tube while in Fig.11b, they are along the length of the tube. This happens because in first case, tubulation occurs due to curved protein while in second case, bundling tendency of the proteins is the driver of tube formation.

**FIG. 11:**
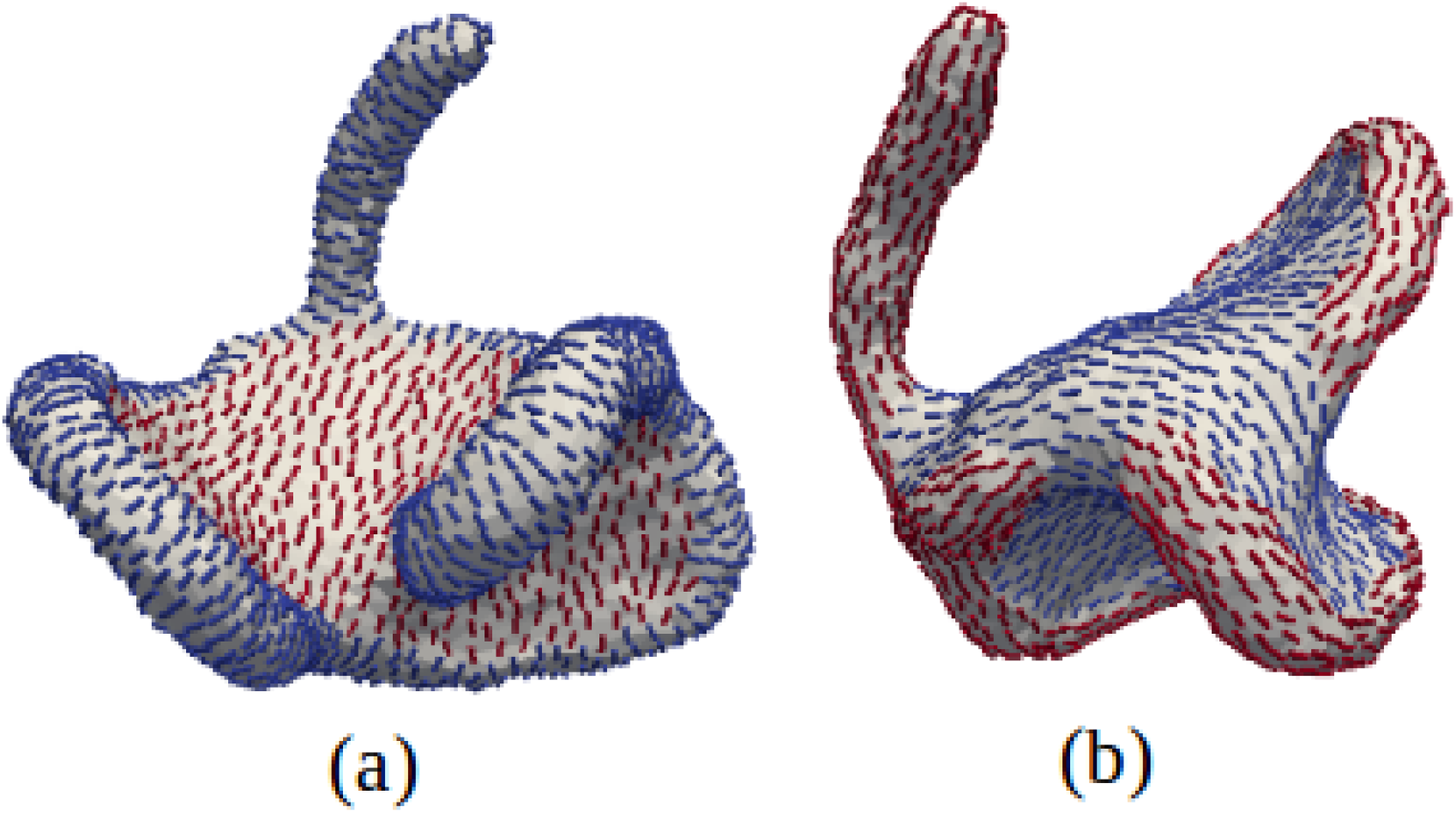
Membrane tubulation due to two different proteins (a) Both proteins curved with curvature value being 1.0 for Blue and −0.05 for Red. (b) One protein is curved with curvature −0.05 (Blue) while the other one is straight *c*_⊥_ = 1.0, *c*_‖_ = 0 and have the tendency to form bundles. Other parameters are *κ* = 20, *κ*_‖_ = 10, *κ*_⊥_ = 10 and 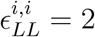 in *k_B_T* unit.

## IV. CONCLUSION

Membrane deformation in cell occurs due to multiple proteins working in synergy with each other. Some processes like clathrin mediated endocytosis^4^ and formation of three way junctions in the endoplasmic reticulum^62,63^ are examples of membrane deformation and stabilisation of induced curvature due to multiple proteins. In this work, we have shown the deformation of membrane due to multiple proteins. These proteins can have different curvatures and interactions. Our simulations show that tubes are forming by positively curved proteins and valley (junction) are formed and stabilized by negatively curved proteins. We also show the effect of interaction between different curved proteins, where we report two particular cases: (i) 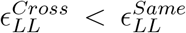 and (ii) 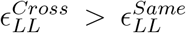. We show that the deformation occurs when 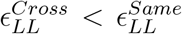 between the two oppositely curved proteins. If the cross interaction is high 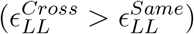, then no deformation occurs because in this case two oppositely curved proteins arrange at random position parallel manner. In this case, proteins are unable to deform the membrane. Further, we show that the membrane tubulation by the mixture of different proteins having a variety of curvatures leading to formation of different morphologies.

Besides the protein and membrane specific parameters, the Hamiltonian can also be augmented to explore features such as effect of surface tension on membrane deformation and protein organization patterns.^39^ To show this ability of the model, we carried out an additional simulation after including an additional term *∫ σ.dA* in the Hamiltonian formulation. Fig.12A-C shows the membrane deformation for three different values of membrane tension *σ* = 5, 10 and 15 (in arbitrary units) where parameters are chosen from Fig.7(*κ*_‖_ = 10 and 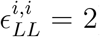). It very clear from the results in Fig.12A-C that higher membrane tension puts a larger penalty on deformation and membrane reticulation.

**FIG. 12:**
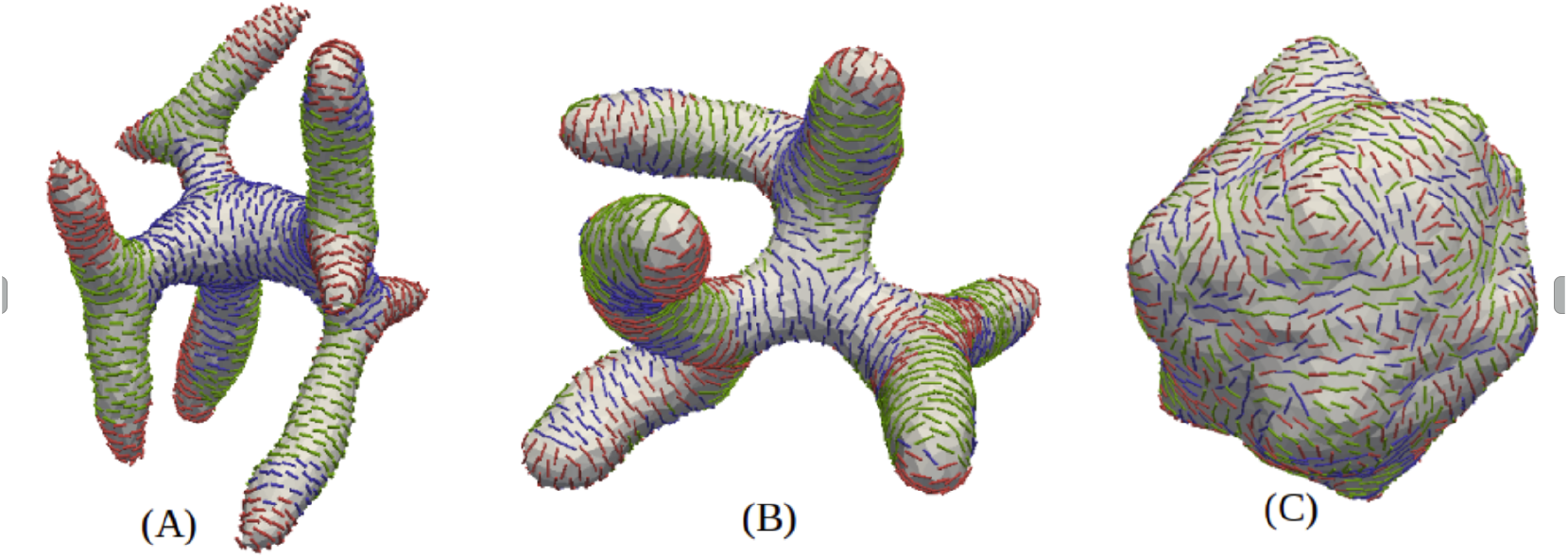
Effect of membrane tension on membrane deformation due to three different proteins.A-C show the deformed shapes of vesicle due to three different values of membrane tension *σ* = 5, 10 and 15 in arbitrary units.Other parameters are *κ* = 20, *κ*_‖_ = 10 and 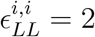 in *k_B_T* unit.

This mesoscopic model can be used as high throughput method to elucidate design-level features that can be used to “engineer” programmed membrane remodelling. Also, it can be successfully used to recapitulate the deformation profile observed in experimental images by scanning and optimizing the parameters space in the augmented Helfrich-based continuum Hamiltonian model. Despite being mesoscopic in nature, the method is able to incorporate the variety and heterogeneity of the biological systems under consideration.

## V. AUTHOR CONTRIBUTIONS

Anand Srivastava conceived the idea and formulated the Hamiltonian with help of Gaurav Kumar. Gaurav Kumar carried out the simulations and all the analyses. Gaurav Kumar and Anand Srivastava wrote the paper.

## VI. ACKNOWLEDGMENTS

Gaurav Kumar would like to acknowledge financial support from DST India, DBT-IISc and SERB. Financial support from the Indian Institute of Science-Bangalore and the high-performance computing facility “Beagle” setup from grants by a partnership between the Department of Biotechnology of India and the Indian Institute of Science (IISc-DBT partnership programme) are greatly acknowledged. A.S. thanks the startup grant provided by the Ministry of Human Resource Development of India and the early career grant from the Department of Science and Technology of India. A.S. also thanks the DST for the National Supercomputing Mission grant. FIST program sponsored by the Department of Science and Technology and UGC, Centre for Advanced Studies and Ministry of Human Resource Development, India is gratefully acknowledged by the authors. This research was also supported in part by the National Science Foundation under Grant No. NSF PHY-1748958 (KITP e-visit).

